# Substrate geometry affects population dynamics in a bacterial biofilm

**DOI:** 10.1101/2023.08.30.555518

**Authors:** Witold Postek, Klaudia Staskiewicz, Elin Lilja, Bartłomiej Wacław

## Abstract

Biofilms inhabit a range of environments, such as dental plaques or soil micropores, often characterized by intricate, non-even surfaces. However, the impact of surface irregularities on the population dynamics of biofilms remains elusive as most biofilm experiments are conducted on flat surfaces. Here, we show that the shape of the surface on which a biofilm grows influences genetic drift and selection within the biofilm. We culture *E. coli* biofilms in micro-wells with an undulating bottom surface and observe the emergence of clonal sectors whose size corresponds to that of the undulations, despite no physical barrier separating different areas of the biofilm. The sectors are remarkably stable over time and do not invade each other; we attribute this stability to the characteristics of the velocity field within the growing biofilm, which hinders mixing and clonal expansion. A microscopically-detailed computer model fully reproduces these findings and highlights the role of mechanical (physical) interactions such as adhesion and friction in microbial evolution. The model also predicts clonal expansion to be severely limited even for clones with a significant growth advantage – a finding which we subsequently confirm experimentally using a mixture of antibiotic-sensitive and antibiotic-resistant mutants in the presence of sub-lethal concentrations of the antibiotic rifampicin. The strong suppression of selection contrasts sharply with the behavior seen in bacterial colonies on agar commonly used to study range expansion and evolution in biofilms. Our results show that biofilm population dynamics can be controlled by patterning the surface, and demonstrate how a better understanding of the physics of bacterial growth can pave the way for new strategies in steering microbial evolution.

## Introduction

Bacterial biofilms are conglomerates of cells bound together by extracellular matrix containing compounds such as polysaccharides and nucleic acids (1). Found widely in natural ecosystems, biofilms play crucial roles in medicine (2, 3) and technology (4, 5). In all these contexts, the emergence of new genetic variants is a concern. For example, cells in a biofilm can acquire resistance to antibiotics through various mechanisms (6, 7), including *de novo* genetic mutations and horizontal gene transfer (8). Limiting the spread of such new variants is desirable, aligning with ongoing efforts to control biological evolution, which are gaining momentum (9).

The growth and population dynamics of biofilms are influenced by biochemical cues (10), competition (11), cooperation (12), cell death (13), and mechanical interactions (14, 15). In particular, the role of cell-cell and cell-surface interactions in the establishment of new variants has been explored in experimental (16, 17) and theoretical (18, 19) studies of colonies growing on agarose gel surfaces. In such colonies, bacteria primarily replicate at the expanding front due to nutrient depletion and waste accumulation at the center (20). When a new mutant arises at the front, it either “surfs” along the advancing front and forms a clonal cluster that expands into a new territory, or it gets outpaced by neighboring clonal populations and loses the competition for nutrients (16, 21-25). The probability of a new variant spreading depends on the interplay of mechanical interactions between bacterial cells and with the substrate, as well as the variant’s fitness compared to the parent strain (13, 20, 26-33). These conclusions hold true also for three-dimensional, thick biofilms, which consist of a growing active layer and a quiescent bulk (34, 35).

A significant limitation in these studies has been the use of flat substrates. In natural environments, biofilms often grow on non-flat surfaces such as rocks and underwater stones (36), pipelines (4), mineralized surface of urinary catheters (37), pores of the human skin (38), porous beads in water treatment plants (39), marine snow (40), and colon crypts (41). Recent work on bacterial colonies growing on rough agar have shown that selection decreases and neutral drift increases compared to flat agar (42). A similar effect has been obtained by placing obstacles in the path of an expanding colony (43). Clonal dynamics is also affected in colonies encountering physical objects during expansion (44). However, these studies primarily focus on quasi-two-dimensional colonies that grow parallel to the rough surface, which does not consider the perpendicular growth of 3d biofilms. Consequently, there is a lack of experimental and *in silico* research on the influence of surface irregularities on the biofilm population dynamics.

To bridge this gap, we employ a microfluidics-based model system to investigate a more realistic scenario of a biofilm growing on a rough surface, where the top of the biofilm is sheared off by flow. We aim to understand how the population dynamics of genetic variants is influenced by substrate roughness. Specifically, we grow biofilms initiated with a mixture of fluorescently labelled cells in microscopic wells, featuring sine-like undulations of the bottom surface. We show that a moderate level of surface corrugation at the scale slightly larger than the cell size reduces both genetic drift and selection strength, enabling coexistence of clones with significantly different growth rates. This holds true even when the amplitude of substrate roughness is much smaller compared to biofilm thickness. We attribute this effect to cell-surface interactions that influence cell orientation and movement in the biofilms. The resulting velocity field hinders clonal mixing everywhere in the biofilm except close to the substrate, where bacteria can orientate and move parallel to the surface. Substrate roughness strongly limits this movement, resulting in narrow clonal sectors that do not invade each other even when clones have different fitness. Our microfluidic experiments, coupled with mathematical modeling, underscore the role of mechanical interactions in biofilm growth and evolution, and suggest a simple and robust way of controlling biofilm genetic diversity by varying substrate roughness.

## Results

### Substrate roughness leads to clonal sectors mirroring the substrate geometry

We cultured biofilms using a microfluidic device (Fig. 1) comprising multiple micro-wells connected to a deeper main channel for delivering growth medium and bacteria to the wells, similar to previously published devices (45, 46). However, in contrast to those studies, each well (100×100×7 μm) had a sine-patterned bottom surface (Fig. 1A), with varying amplitude and period across different wells, including some with flat bottoms. The amplitude of surface height modulation was only a small fraction of the total well height, and the sine-like indentations did not physically isolate different regions of the biofilm. The quasi-two-dimensional well shape allowed for the formation of multi-layered biofilms similar to naturally occurring biofilms, but the biofilms remained thin enough for cell movement to be limited mostly to the XY plane. This facilitated imaging of the biofilm using a wide-field epi-fluorescent microscope.

**Fig. 1.**
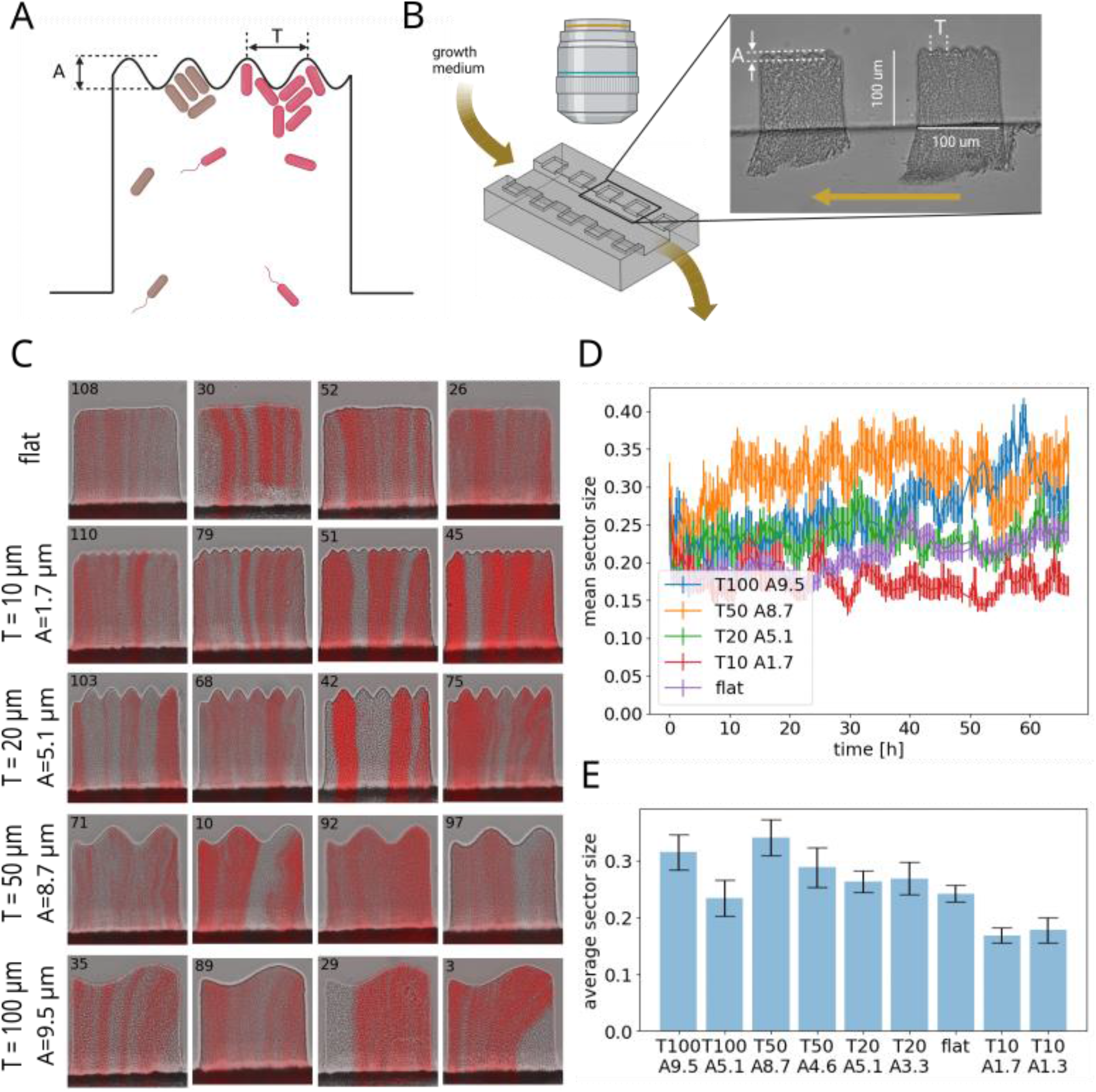
Substrate geometry affects the number and size of clonal sectors. **(A)** Illustration of a single 100×100 μm well with a corrugated bottom for biofilm growth. **(B)** A simplified diagram of the experimental set-up. The actual microfluidic device has 240 wells with sine-like corrugations of 8 different amplitudes *A* and periods *T*, each replicated 20 times, along with 80 flat-bottomed wells. **(C)** End-point snapshots of randomly selected wells with different configurations. The horizontal extension of quasi-vertical clonal sectors often matches the surface undulations. **(D)** Mean sector size versus time for five different combinations (*A, T*). *t* = 0 h is 4 days post-inoculation. During the initial 4 days some biofilms fell out and were re-established. **(E)** Mean sector size as fraction of the well’s width at the end point of live imaging (t=65 h, total growth time 6d 15h) for all well types (all combinations (*A, T*)). Error bars are S.E.M.

To examine the impact of growth on corrugated surfaces on clonal dynamics within the biofilm, we performed standing variation experiments using the biofilm-forming strain *E. coli* 83972 and its derivatives (*Methods*). We inoculated the microfluidic device with a 1:1 mixture of similarly-fit (*SI Methods*) red fluorescent (mKate) and non-fluorescent cells. Nutrient broth (LB) was pumped through the main channel, while we monitored all wells using fluorescence and phase-contrast microscopy. The experiment was conducted at room temperature (24-26°C) to reduce growth and clogging in the main channel. Nutrient flow trimmed excess biofilm extending into the main channel, thus allowing nutrients to diffuse into the wells. Biofilms quickly formed in all wells, with initial patches of fluorescent and non-fluorescent bacteria developing into sectors aligned with the direction of biofilm growth. The number of sectors and their positions matched the undulating pattern of the well bottom (Fig. 1C and SI Video 1), especially evident for undulations with periods much smaller than the well’s width. Most sectors were parallel to the side walls, but some exhibited bending (e.g., biofilm no. 3 in Fig. 1C), suggesting non-uniformities in the velocity field within the well. Once established, the sectors maintained nearly constant width during live imaging (Fig. 1D).

To quantify the difference between sector patterns in various wells, we calculated the mean sector size as a fraction of the well’s width (*Methods* and *SI Methods*). Figure 1D shows that long-period undulations result in larger sectors compared to flat-bottomed and short-period ones. To assess the impact of the amplitude *A* alone, we compared wells with two different amplitudes but identical *T*. Figure 1E shows that reducing the amplitude has a lesser effect on the size of neutral sectors compared to decreasing the period *T* for small *T*.

In contrast to similar experiments in bacterial colonies (16, 18, 21, 24, 25), the fluorescence within the sectors appeared more heterogeneous, indicating that the sectors were not entirely monoclonal. To understand how genetic diversity within each sector was reduced due to clonal expansion in the pockets of the undulating surface, we analyzed the inter-pocket diversity and compared it with intra-pocket diversity (SI Fig. S2). Specifically, we calculated the standard deviation of fluorescence intensity within each pocket (SI Fig. S2A). We then calculated the ratio *ρ* of the average standard deviation of within-pocket fluorescence and the standard deviation of mean within-pocket fluorescence. SI Figure S2B shows that *ρ* decreases with decreasing pocket size (decreasing *T*). Additionally, for a given *T*, the ratio *ρ* is smaller for undulated surfaces than for flat-bottom wells when computed in “virtual pockets” positioned as they would be in the corrugated well. This indicates that while both fluorescent and non-fluorescent bacteria may be present in some pockets, undulations significantly hinder the mixing of adjacent bacterial populations.

### Bent sectors arise from the non-uniformity of the velocity field caused by heterogeneous adhesion

Some fluorescent sectors in Fig. 1C are bent, which suggest non-uniform flows in the biofilm (see also SI Video 1). This non-uniformity could be a result of differences in the growth rate or variations in adhesion to the surface in different parts of the biofilm. To distinguish between these two possibilities, we calculated the velocity field in the biofilm from short bright-field time-lapse videos recorded at a higher frame rate (1 fps) than those used for Fig. 1, using an optical-flow based method (*Methods* and *SI Methods*). We then calculated the local growth rate *g*(*x, y*) using the continuity equation for an incompressible fluid with local mass generation: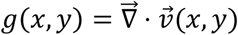. Figure 2A shows the velocity field superimposed on the field representing the growth rate for selected wells. It is evident that, except for a small gradient towards the main channel and the direction of growth (the vertical direction in the picture), the growth rate remains relatively uniform, even though the velocity field occasionally has a significant horizontal (perpendicular to the growth direction) component. This suggests that most of the observed heterogeneities of the velocity field are due to stronger adherence of parts of the biofilm to the sides of the wells.

**Fig. 2.**
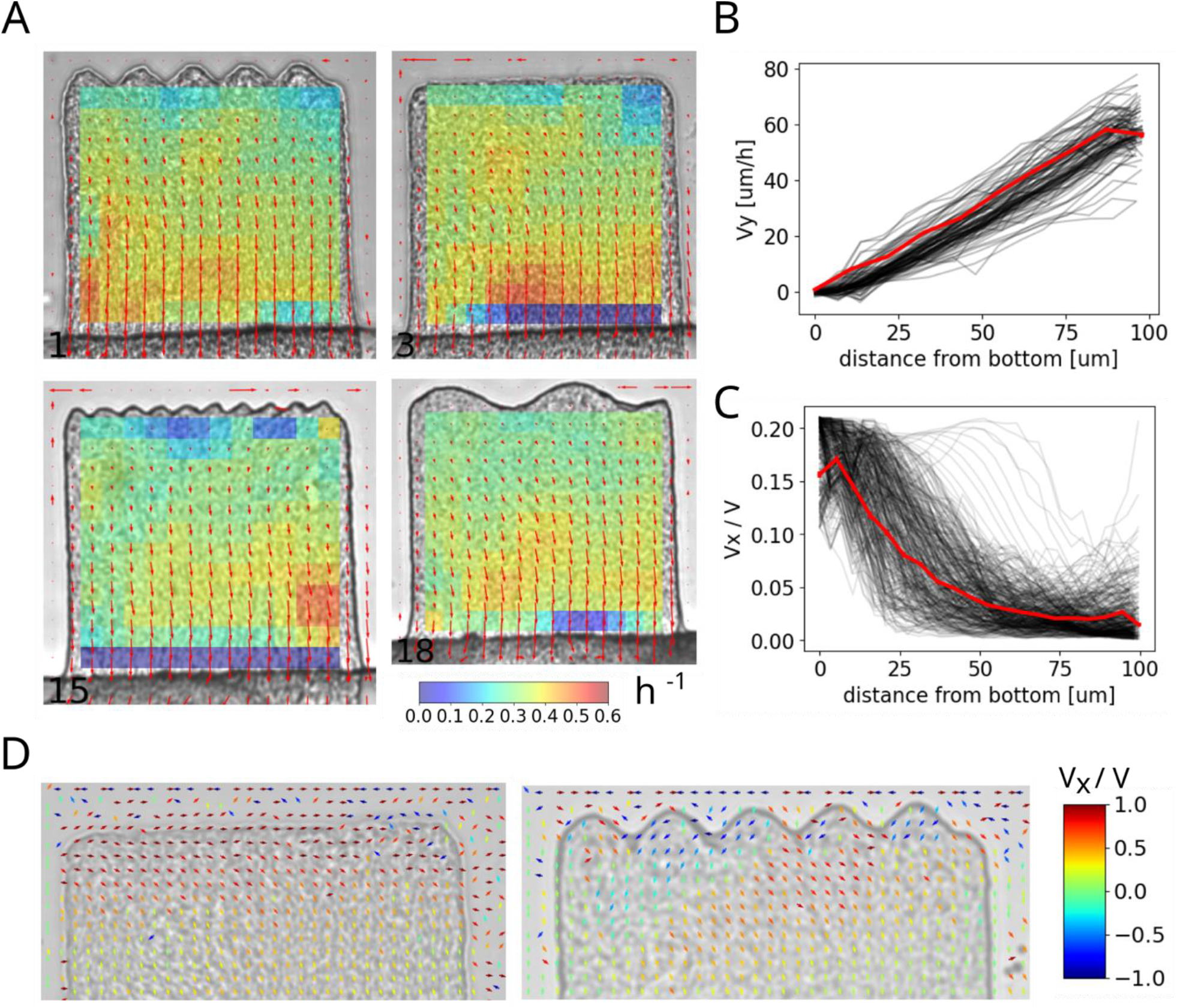
Velocity field in the wells. **(A)** Examples of the velocity field (arrows) and growth rate (color map) for wells of different types. **(B)** Average vertical velocity (red curve) as a function of distance from the bottom surface of flat-bottomed wells. The individual black lines represent velocities measured at various horizontal positions (approx. 10 per well) within the well, and across different wells. **(C)** Average horizontal component of the velocity field (red curve) versus the distance from the well bottom. The black lines are as in panel C. **(D)** Zoomed-in plots of the velocity field near the bottom for a corrugated- and a flat-bottom well. Arrows are color-coded based on the *V*_*x*_/*V* component of the field (scale bar).

### Spatial isolation arises from the focusing property of the velocity field

Next, we investigated the properties of the velocity field that could explain the spatial isolation of the sectors. Figure 2B shows the vertical component of the velocity field, averaged across all horizontal positions and all flat-bottom wells. The linear increase of the vertical velocity with distance from the bottom indicates that the growth rate in our system is the same at all depths inside the biofilm. Such a linear velocity field facilitates the orientation of rod-shaped cells in the direction of growth (47). Figure 2C shows that the horizontal component of the velocity field, plotted as a fraction of the total velocity, decreases rapidly with distance from the bottom. Consequently, significant horizontal cell movement occurs primarily near the bottom surface of the well. This movement enhances genetic drift, promoting the establishment of a single clone in flat-bottom wells. To assess whether the presence of surface undulations hinders this movement, we examined the velocity field near the bottom of flat- and corrugated wells. Figure 2D shows that the horizontal movement of cells is indeed disrupted by the presence of surface undulations.

### Computer model reproduces the experimental results

To understand the experimental results, we used a 2d computer model similar to the one reported before (20), to simulate a colony of rod-shaped bacteria in a square well with one open side, through which cells could escape and be removed from the system. We assumed that all bacteria in the biofilm divided at equal rates as supported by Fig. 2B. To facilitate tracking clonal sectors, the system was initialized with bacteria each assigned a different, heritable genotype, represented by a random color. The simulation ran for 72 h (a similar duration as the experiment) for different pairs of *A, T*. Figure 3A shows that, as expected, undulated bottom surface constrained the spread of genetic variants to sectors originating from within each sine-like pocket. The number of sectors decreased in time but eventually stabilized for corrugated surfaces (Fig. 3B). The final number of sectors closely aligned with the number of undulations (Fig. 3C, D), for a broad range of amplitudes *A* (Fig. 3E). In contrast, a flat- or only slightly undulated bottom surface allowed bacteria to spread along it, leading to increased genetic drift. Figure 3F, which is analogous to the experimental figure 2C, shows that horizontal movement was indeed limited near the bottom surface. A new clonal variant could only establish if initiated by a cell close to the substrate on which the biofilm grows (Fig. 3G), otherwise the variant would be pushed out of the well by replicating cells that were closer to the substrate. This is the opposite of what occurs in bacterial colonies growing on a flat agar surface, where new variants can only establish if they emerge close to the colony edge (18, 20).

**Fig. 3.**
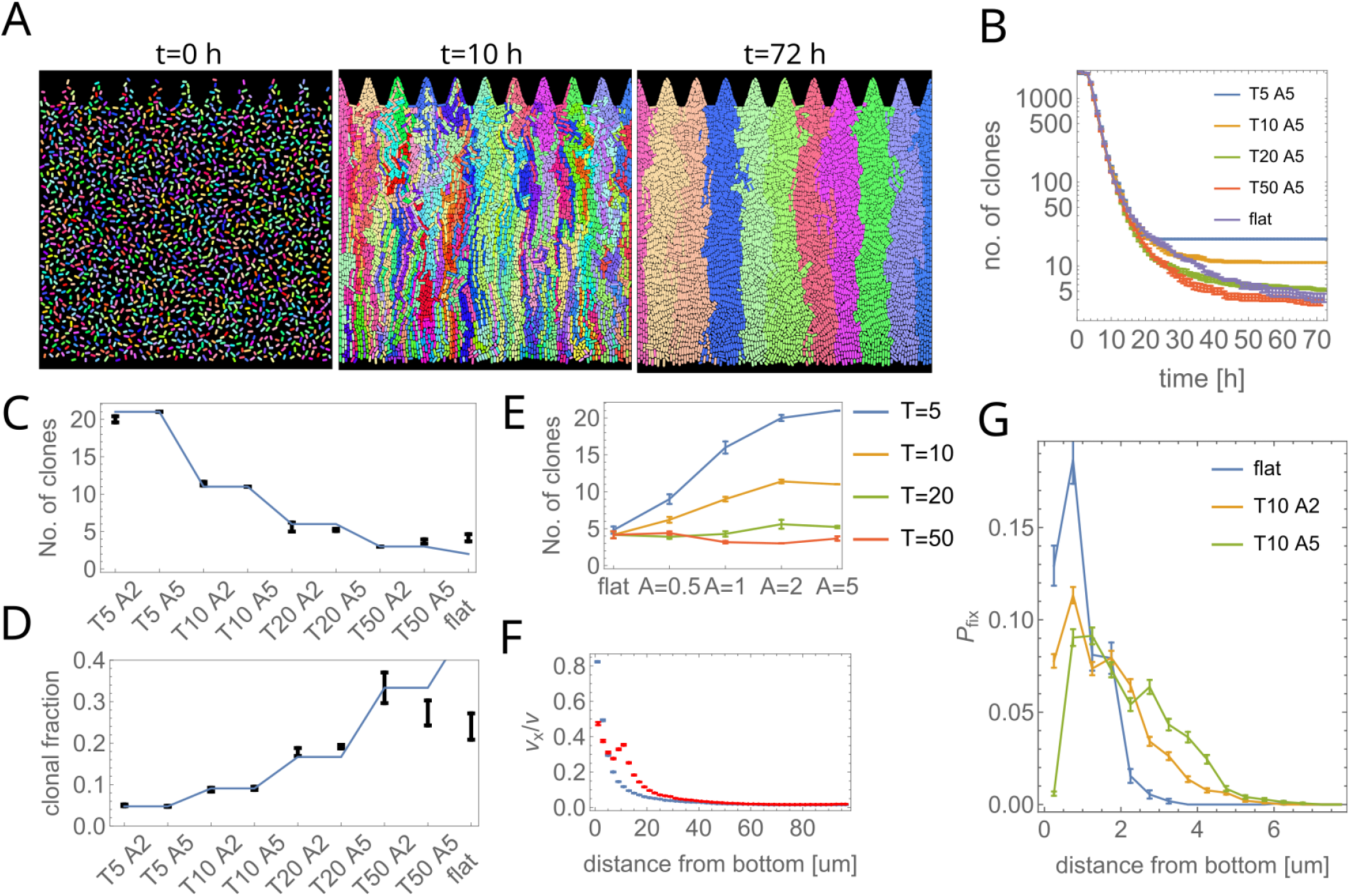
Computer model of the experiment. **(A)** Simulation snapshots for *T* = 10, *A* = 5. **(B)** The number of clones as a function of time, for different *T, A*. **(C, D)** The number of clones after *t* = 72 h (C) and the mean fraction of the population occupied by a clone (D), for different *T, A*. The blue line shows theoretical average values for the number of clones *N*_clones_ = (*T* + 100 μm)/*T* and the fraction occupied by a single clone *f* = 1/*N*_clones_ based the number of pockets, under the assumption of intra-pocket clonality. **(E)** The number of clones (as in panel C) as a function of amplitude *A*, for different periods *T*. **(F)** The lateral component of the local velocity field as a function of the distance from the bottom. The blue curve is for a flat bottom well, the red curve is for *T* = 10, *A* = 5. **(G)** Probability density function that the progeny of a cell, originated at *t* = 2 h at a distance *y* from the bottom, survives until *t* = 72 h. In all panels, error bars = S.E.M.

### Model prediction that undulations serve as suppressors of selection is confirmed experimentally

So far, we considered neutral variants, i.e., all genotypes (both in the experiment and in the model) had nearly identical growth rates. Since spatial patterning of the bottom surface decreases interactions between neighboring populations of bacteria, we reasoned that it should also prevent fitter variants from taking over. To test this hypothesis, we initialized our computer model with a 10:1 mixture of normal-growing: faster-growing clones with relative fitness *W* = 1.5 (*Methods*), and measured the fraction of the less-fit clone after *t* = 72 h. Figure 4A shows that the less-fit clone is quickly outcompeted by the faster-growing variant in flat-bottomed wells, but undulations impede its spreading, resulting in a substantial fraction (40-60%) of the less-fit clone remaining (Fig. 4B). Moreover, when simulated bacteria adhere to the surface of the well, the effect of selection is further diminished (dark green points in Fig. 4B). Therefore, our system acts as a suppressor of selection (48), allowing coexistence of diverse genetic variants.

**Fig. 4.**
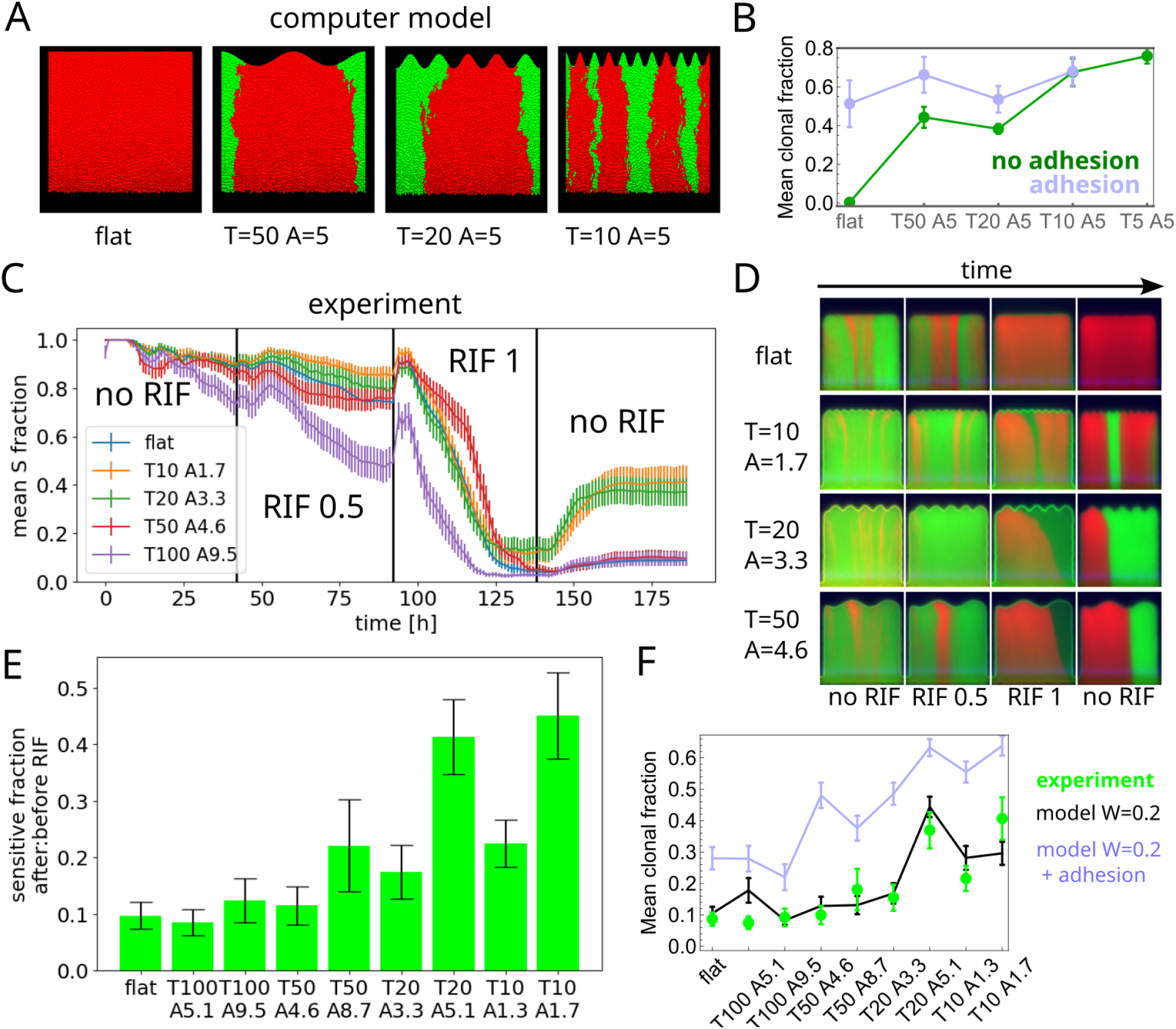
Corrugated surface limits selection. **(A)** Computer simulation snapshots (*t* = 72 h) illustrate how corrugations constrain the spread of the fitter mutant (red, relative fitness *W*_*R*/*G*_ = 1.5). **(B)** The fraction of the less-fit green strain remaining in the biofilm after 72 h increases as the corrugation period *T* decreases, for absent or weak adhesion. Strong adhesion nullifies this effect. **(C)** Experimental validation of the model: the fraction of RIF-sensitive (less fit) green strain as a function of time in an experiment in which the strength of selection was varied in time by adjusting the RIF concentration. The fitter red strain dominated in flat-bottomed and large-*T* wells after transient RIF exposure, while rapid undulations (small *T*) significantly limited the spread of the fitter mutant. **(D)** Snapshots of wells corresponding to different phases of the experiment from panel C. **(E)** The ratio of the sensitive strain fractions at two time points: after (*t* = 182 h) and before (*t* = 41 h) the exposure to RIF. **(F)** The mean fraction of the sensitive strain in different types of wells (green points) is reproduced by the computer model without adhesion (black line). Error bars are S.E.M.

To experimentally test this prediction, we inoculated our microfluidic device with a 1:10 mixture of mKate RIF^R^ red-fluorescent, rifampicin (RIF) resistant strain and GFP RIF^S^ green-fluorescent rifampicin sensitive strain. In the absence of RIF, the red strain had a small fitness disadvantage (*W*_*R*/*G*_ ≈ 0.9) compared to the green strain (SI Fig. S5). By varying the RIF concentration, we could thus tune the selective advantage of the red versus green strain. After approx. 40 h of incubation in pure LB medium in the microfluidic device, which resulted in the establishment of fluorescent sectors, we changed the medium to 0.5 μg/ml RIF in LB, and then after another 50 h to 1 μg/ml RIF. Figure 4C shows that the fraction of the RIF-sensitive green fluorescent strain decreased in all wells during the exposure to RIF, with a more pronounced reduction in wells with flat bottoms, and corrugated bottoms with wider undulations. Subsequently, we replaced RIF with pure LB medium, leading to the successful re-establishment of the sensitive strain in corrugated wells with *T* < 100 μ*m* (Fig. 4C, D, and SI Video 2). However, re-establishment occurred only rarely in flat-bottom wells or those with *T* = 100 μ*m*. Figure 4E quantifies this effect by plotting the ratio of the sensitive strain fractions at two time points: after (t=182 h) and before (t=41 h) the exposure to RIF, for different well types. A similar trend is visible in the number of RIF-sensitive sectors (SI Figs. S3 ad S6).

The rate with which a fitter sector expands can be related to the relative fitness of the green versus the red strain (*SI Methods*). By fitting the model to the experimental time-series data for flat-bottom wells, we determined the relative fitness *W*_*G*/*R*_ ≈ 0.2 (SI Fig. S4). We then ran the computer model without adhesion using initial clonal fractions obtained at the end of the 0.5 μg/ml RIF phase of the experiment from Fig. 4C, the relative fitness *W*_*G*/*R*_ = 0.2 and undulations amplitudes as in Fig. 4E. The model correctly reproduced the experimental results (Fig. 4F). Interestingly, repeating the same procedure for the model with adhesion sufficiently strong to affect the velocity field of the growing biofilm yielded a much worse fit (Fig. 4F). This does not mean that cells did not adhere to the walls but only that surface undulations were more important than adhesion for the outcome of our experiment.

## Discussion

We have shown that biofilm population dynamics depends on the shape of the surface on which the biofilm grows. By transitioning from a flat to a corrugated surface, we have demonstrated that surface undulations of appropriate period and amplitude can prevent clonal sub-populations from invading each other. This reduces genetic drift, enabling neutral variants to coexist, while also diminishing the impact of selection, preventing fitter variants from dominating over less-fit clones. These effects arise from the interplay of bacteria-surface interactions that influence cell orientation and the velocity field in the biofilm. Importantly, the depth of the undulations required to see these effects is only a small fraction of the biofilm’s thickness, and the undulations do not physically separate different regions of the biofilm.

Our model system demonstrates significant differences compared to bacterial range expansion experiments on agar plates (21, 24, 25, 43, 49). Clonal sectors emerge despite uniform growth throughout the entire biofilm volume, whereas on agar plates a thin growing layer is necessary for clonal segregation to occur. Additionally, in our system, the roughness of the biofilm’s top surface has no effect on its population dynamics, unlike in colonies on agar where the roughness of the leading edge significantly affects genetic drift (20). This is because in our system it is the growth at the bottom, not at the leading edge (top surface), that drives the dynamics of clonal sectors. Interestingly, “static” i.e., time-independent undulations of the bottom surface have the opposite effect compared to dynamically changing undulations of the expanding frontier of a growing colony (16, 43): genetic drift is reduced rather than enhanced by “static” roughness.

Computer simulations from Fig. 4 show that surface adhesion could have a profound effect on selection, reducing the influence of growth rate differences on the probability of establishment of fitter variants. Since in our model fitter clones must first expand close to the bottom surface to dominate the biofilm, we conclude that adhesion could partially counterbalance the effect of increased growth rate and hinder expansion. In our experiment, this effect does not seem to be very strong, perhaps because inter-cellular adhesion is stronger than adhesion to the walls. Nevertheless, we hypothesize that mutants with reduced adhesion could gain an additional, growth-independent selective advantage, similarly to what happens in bacterial colonies (50). However, the advantage might be short-lived: since adhesion is essential for biofilm establishment on surfaces, less-adherent variants could eventually cause the biofilm to detach. Consequently, conducting an experiment with a less adherent strain of bacteria would not be possible in our setup as it would result in biofilms falling out of the wells.

Surface adhesion is likely responsible for the observed persistence of neutral sectors in some flat-bottomed wells. In a well-mixed growing population whose size remains approximately constant due to the continuous removal of surplus population, one clone would eventually always reach fixation (51). However, in our system, adhesion may prevent fixation, particularly if certain cells (e.g., older ones) adhere stronger than others. Such cells may persist in the well for an extended period, leading to the formation of long-lasting sectors by their progeny.

A key mechanism driving clonal expansion on flat surfaces and within sine-like pockets is the horizontal component of the velocity field near the surface (Fig. 2). In the computer model (Fig. 3), this component arises due to “buckling”, i.e., pressure-driven bending of chains of bacteria experiencing growth-induced compression (28, 30, 52, 53). This is supported by the lack of lateral motion of cells in simulations in which the height of the biofilm has been made much smaller, reducing mechanical stress and making cells more aligned with the flow, and the restoration of lateral movement upon increasing the friction coefficient (SI Videos 3, 4, 5). In the actual experiment, buckling manifests as bends in fluorescent sectors, particularly noticeable near the bottom. Notably, buckling is essential for the velocity flow to acquire a horizontal component, causing cells at the bottom of the well to orient themselves perpendicularly to the surface. For undulated surfaces, the onset of lateral buckling occurs earlier i.e., for smaller compression forces, because the curvature of such surfaces prevents cells from aligning into chains. This intuitive explanation clarifies why undulating surfaces hinder mixing: earlier buckling (more pronounced for surface undulations of higher amplitude) results in a faster transition from horizontal to vertical biofilm flow.

Since buckling is affected by adhesion and friction not only with the bottom and side walls but also the glass and PDMS surfaces, we expect that our results would be quantitatively – but not qualitatively – different, if we used taller wells (*h* > 10 μm) to accommodate more cell layers. Creating a fully three-dimensional biofilm would reduce the role of such boundary effects. Patterning the surface in two directions would be necessary to limit clonal expansion under such conditions.

We speculate that our findings generalize to naturally occurring biofilms that are relatively thin (a few hundred μm) to enable growth at the bottom. If the biofilm is thick enough to prohibit cell division at the bottom but its height remains limited due to mechanical shearing or flow, the shape of the surface it adheres to should become less important. Nevertheless, this might change in the presence of particles such as microplastics attaching to the biofilm and creating new adhesion points for cells.

Our results suggest that bacterial population dynamics in the biofilm can be controlled by patterning the surface. Many population genetics models (48, 54-58) rely on compartmentalization to influence population dynamics, which may not be applicable to growing biofilms. In contrast, our approach leverages the physics of the biofilm to achieve effective spatial separation. By manipulating the surface geometry, we can restrict the expansion of undesired genetic variants; we demonstrate this specifically for an antibiotic-resistant mutant. Sub-MIC transient antibiotic exposure is not uncommon and may lead to the emergence of resistance (46, 59). However, further research is required to see if surface patterning could be used for medical devices such as catheters or implants, which are prone to biofilm invasion (37). Similar strategies could also be utilized to stabilize engineered bacterial communities (60), in particular for the use in biosensing (61) or bioremediation, in which different bacterial ecotypes must often coexist together (62), and mutations in synthetic gene networks often present in such communities must be prohibited.

## Materials and Methods

### Microfluidic device

We fabricated a two-layer device made of PDMS attached to a glass slide (SI Fig. S1), with 240 micro-wells on both sides of a 500 μm-wide and 87 μm-deep channel. Each well measured 100×100×7 μm (width (X) x height (Y) x depth (Z)). The device contained 20 replicates of each of 8 sine wave/amplitude combinations, and 80 flat-bottomed wells. To fabricate the device, we utilized soft lithography, following well-established protocols (*SI Methods*). A photomask designed in AutoCAD (Autodesk) and printed by MicroLitho, UK, was used to expose a layer of negative photoresist on a silicon wafer. After developing, the negative mold was covered with PDMS (Sylgard, Dow Corning) mixed at a 1:10 ratio of curing agent to monomer and baked at 75°C for at least four hours. The resulting PDMS device was peeled off, oxygen-plasma treated, and bonded to a plasma-treated 1 mm-thick glass slide. After inserting PTFE tubing (Bola Bohlender, Germany, I.D. = 0.5 mm, O.D. = 1.0 mm), the device was connected to syringe pumps PHD2000 (Harvard Apparatus, USA) or (in some experiments) SyringeONE Programmable Syringe Pump (Darwin Microfluidics). We used plastic syringes with appropriate media (bacterial culture, LB, LB + rifampicin), depending on the type of experiment. Prior to use, all devices were flushed with 70% ethanol (Figs. 1-2) and 70% ethanol + 5% NaOH (Fig. 4) to sterilize the device, remove air and enhance bacterial adhesion.

### Bacterial strains

We used the *E. coli* strain 83972 (63) and its fluorescent derivatives for all experiments, with either mKate (red) or GFP (green) being constitutively expressed from the bacterial chromosome. For the experiments with a RIF-resistant strain we used a variant of the red fluorescent strain with resistance conferred by a single point mutation of the *rpoB* gene. All strains easily adhered to surfaces and formed biofilms. The presence of the fluorescent reporter did not reduce fitness compared to the ancestral strain, but rifampicin resistance decreased fitness by about 10% (SI Fig. S5). All genetic modifications are detailed in *SI Methods*.

### Biofilm growth and imaging

All experiments were performed at room temperature (24-26°C). We inoculated the microfluidic device through the attached PTFE tubing using dense bacterial cultures that were grown overnight in LB (Miller) broth (Carl Roth, Germany) at 37°C/180 rpm, diluted in fresh LB, mixed in desired ratios (1:1 or 1:10 as established by OD_600_ measurements) and centrifuged to increase the cell concentration. After allowing bacteria to settle in the wells for approx. 30 min, we swapped the syringe with the bacteria to a new one filled with LB broth and initiated the flow. We used variable-flow rate protocols (*SI Methods*) to reduce clogging and biofilm growth in the main channel. For the experiment in Fig. 4, we replaced the LB medium with LB + RIF, and later switched back to LB, as described in the main text.

### Microscopy and image analysis

Images were acquired on two fully automated Nikon Ti2-Eclipse epi-fluorescent microscopes equipped with automated XY stages, the Perfect Focus System, GFP and mCherry fluorescent filters, and controlled by MicroManager (64). Depending on the experiment, we used either 20x or 40x long-working distance Nikon objectives. Custom-written Python and Mathematica® code was used to load and process raw TIFF images outputted by MicroManager, and to analyze and plot the data. See *SI Methods* for details.

### Computer Model

Computer simulations were conducted on the Edinburgh compute cluster at SoPA. We adapted the model previously reported (20), which involved representing bacteria as growing spherocylinders, with their movement confined to the XY plane, and interacting mechanically through Hertzian-like repulsion. The dynamics was overdamped with Stokes-like friction proportional to the velocity of the moving cell. We did not model nutrient diffusion and assumed that all cells of the same type grew at the same rate, regardless of their location in the well. We modelled repulsion from the walls of the well as a force that increased linearly with the overlap between the cell and the wall, with a stiffness constant 10^7^ pN/μm sufficiently large to prevent the overlap to increase beyond a fraction of a μm. Adhesion between bacteria and walls was represented using elastic springs (spring constant *k* = 10^6^ pN/μm), connecting the center line of the cell to the closest point of first contact on the wall. The springs broke when extended by more than 20% (Fig. 4B) and 5% (Fig. 4F) of their initial length.

## Supporting information

SI Video 1 - biofilm growth in selected wells (as in Fig. 1) over the course of 65 h.

SI Video 2 - biofilm growth in 4 wells of different type: flat, (T,A)=(10,1.7), (T,A)=(20,5.1), and (T,A)=(50,8.7).

SI Video 3 - simulated biofilm in an 80×70 um well, for the same parameters as simulations presented in Fig. 4.

SI Video 4 - simulated biofilm in an 80×30 um well, for the same parameters as SI Video 3.

SI Video 5 - simulated biofilm in an 80×30 um well, for the same parameters as SI Video 3 except for a 20x larger friction coefficient

## Data, materials, and software availability

All code (C++, Jupyter notebooks, Mathematica notebooks), processed image data and simulation results are available at https://github.com/Dioscuri-Centre/biofilms_on_corrugated_surfaces. Due to large file sizes (several TBs), raw image data have not been uploaded. Access to such data will be provided on request.

## Acknowledgements

This work was supported by the following grants: the Bekker Programme of the Polish National Agency for Academic Exchange (NAWA) grant no. PPN/BEK/2020/1/00333/U/00001 (W.P.), the START scholarship of the Foundation for Polish Science (FNP) grant no. 069.2021 (W.P), NAWA Polish Returns grant no. PPN/PPO/2019/1/ 00030 /U/0001 (W.P. and B.W.), and Dioscuri, a programme initiated by the Max Planck Society, jointly managed with the National Science Centre in Poland, and mutually funded by Polish Ministry of Science and Higher Education and German Federal Ministry of Education and Research, grant no. UMO-2019/02/H/NZ6/00003 (W.P., K. S., E. L., B. W.). Figure 1 was created with BioRender.com.

## Competing interests

The authors do not report any competing interests.

## Use of AI tools

ChatGPT was used to refine the text and improve its flow, although no ChatGPT-generated sentences were copied verbatim into the manuscript. The manuscript was read and approved by all authors.

## Notes

https://github.com/Dioscuri-Centre/biofilms_on_corrugated_surfaces

### Competing Interest Statement

The authors have declared no competing interest.

