## Supplementary Methods and Figures for "Substrate geometry affects population dynamics in a bacterial biofilm"

*Microfluidic device architecture*

The microfluidic chip used in our study was a two-layer device consisting of a PDMS mold attached to a 1 mm-thick glass slide (SI Fig. S1). The device featured one inlet and two outlets equipped with pillar-based filters to prevent inflowing debris. The main channel was 500 µm wide. Micro-wells for culturing biofilms extended perpendicularly from the main channel; each was 100 µm wide (parallel to the main channel) and 100 µm deep (perpendicularly to the long axis of the main channel). The final height of the main channels and the wells obtained through soft lithography (see below) was 87 and 7 µm, respectively, so that only a few layers of cells could fit into each well, whereas the much taller main channel allowed for rapid nutrient medium flow. The 7 um well thickness facilitated optical imaging while at the same time ensuring that most bacteria interacted with other bacteria in the bulk of the biofilm rather than with the top (PDMS) or bottom (glass) surfaces. The biofilms were continuously trimmed by the flow in the main channel, ensuring a steady supply of nutrients to the deepest layers of the biofilm.

Each microfluidic device had 240 micro-wells, with 120 on each side of the main channel. The bottom of the well, which faced away from the main channel, was designed to be either flat or undulated. The shape of the undulations was a sine function of different periods and amplitudes (all dimensions in µm): $(T, A) = ((100,9.5),(100,5.1),(50,8.7),(50,4.6),(20,5.1),(20,3.3),(10,1.7),(10,1.3))$. Due to the limitations of soft lithography and mask resolution, it was not possible to have the same amplitude $A$ for all periods $T$. The reported amplitudes are actual amplitudes obtained from microphotographs of the device. Each device contained 20 replicates of each sine wave / amplitude combination, as well as 80 flat-bottomed wells. The CAD design of the device is available on GitHub (1).

*Soft lithography*

To create a negative of the microfluidic device, we followed well-established photolithography protocols (2). First, we designed a photomask in AutoCAD (AutoDesk) and had this mask printed by an external company (MicroLitho, UK). We then covered a 3-inch silicon wafer (Microchemicals) with an SU-8 photoresist (Kayaku Advanced Materials) using a spin coater (Laurell, USA). After a soft bake on a programmable hot plate (4 minutes at 95°C), we exposed the wafer through the photomask representing the micro-wells layer, using a MJB4 mask aligner (SÜSS MicroTec). The second layer of SU-8 was then spun on the wafer, and the wafer was again soft-baked (5 minutes 65°C, 20 minutes 95°C). Edge bead-removal procedure was applied using the spin coater by covering the edge of the spinning wafer with photoresist developer mr-600 (Micro Resist Technology, Germany), deposited through a syringe, to unravel the alignment marks from the first layer. The wafer then underwent another soft bake (5 minutes 65°C, 20 minutes 95°C). Next, the wafer was illuminated through the photomask representing the main channel, and a post-exposure bake (5 minutes 65°C, 10 minutes 95°C) was performed. The specific spinning times, speeds, soft bake/hard bake times, and illumination parameters were obtained from the SU-8 manufacturer's (Kayaku) protocols. The wafer was developed with mr-600 developer (Micro Resist Technology, Germany) according to the manufacturer's protocol. After the development, the wafer was hard baked at 150°C for 30 minutes.

*Microfluidic device casting*
The wafer with the photo-resist negative of the device was covered with PDMS (Sylgard, Dow Corning) mixed at a 1:10 ratio of curing agent to monomer, and baked at 75°C for at least four hours. The cured PDMS was peeled off the wafer and inlet and outlet holes were created using a 1 mm-diameter biopsy puncher (Kai Medical). The PDMS device was activated with oxygen plasma alongside a glass slide in a plasma cleaner (Harrick Plasma, USA) for 60 seconds. Following this, the PMDS mold was gently placed on the glass slide and lightly pressed with a pair of metal tweezers to ensure proper bonding of the PDMS to the glass.

*Bacterial strains*
*E. coli* 83972 (DSM number 103539) was obtained from DSMZ GmbH. The red fluorescent mKate marker (under the control of the constitutive promoter *PtetO1*) was introduced into 83972 using plasmid mediated gene replacement (3, 4) replacing the *galK* gene. The strain used to amplify the mKate marker with the *PtetO1* promoter from was a gift from Meriem El Karoui (5). The green fluorescent GFP marker under the control of the constitutive *PA1* promoter was introduced into *E.coli* 83972 using plasmid mediated gene replacement, replacing the *galK* gene. The GFP marker with the *PA1* promoter was amplified from plasmid pGRG36-Kn_PA1-GFP(6) Plasmid pGRG36-Kn-PA1-GFP was a gift from Frank Rosenzweig (Addgene plasmid # 79088 ; http://n2t.net/addgene:79088 ; RRID:Addgene_79088). The primers used to amplify the upstream and downstream regions surrounding the *galK* gene of 83972, as well as the primers used to amplify the mKate and the GFP markers can be found in Table S1. Crossover PCR was used to anneal the 83972 homologous regions with the mKate and GFP markers, and these constructs were then inserted by restriction digestion and ligation into the plasmid pTOF24 (4) used for the gene replacement.
A rifampicin (RIF) resistant version of 83972 with the mKate marker was generated by plating the strain on LB agar plates supplemented with 100 µg/ml rifampicin, and randomly picking a resistant colony.

| | |
|---|---|
| 1.galK_up_fwd | AAA AAC TGC AGA CAC TGG TTA GCC GTT GTA C |
| 2.galK_up_rev | ATA GGG ACT CGA TTTC TTA CAC TCC GCA TTC |
| 3.mKate_gal_fwd | GGA GTG TAA GAA TCG AGT CCC TAT CAG TGA |
| 4.mKate_gal_rev | CGG GAG TTT CGT TTA TCT GTG CCC CAG TTT |
| 5.galK_down_fwd | GGG CAC AGA TAA ACG AAA CTC CCG CAC TGG |
| 6.galK_down_rev | AAA AAG TCG ACT GAT CGC CAT CAT CTG AAC T |
| 7.Gal_up_fwd | AAA AAC TGC AGT GAC GAT CGT TCT GGT TCA C |
| 8.Gal_PA1_up_rev | TGA TAA CCG CTA CGG AAG AGC TGG TGC CTG |
| 9.PA1_gal_fwd | CCA GCT CTT CCG TAG CGG TTA TCA AAA AGA |
| 10.GFPt7_gal_rev | GGA GTG TAA GAA TCA GCA AAA AAC CCC TCA |
| 11.Gal_GFPt7_down_fwd | GTT TTT TGC TGA TTC TTA CAC TCC GGA TTC |
| 12.Gal_down_rev | AAA AAG TCG ACA CAC TGG TTA GCC GTT GTA C |

Table S1. Primers used to amplify the upstream (1,2) and downstream (5,6) regions of the galK gene of 83972, with overlap to PtetO1_mKate, primers used to amplify mKate with the PtetO1 promoter (3,4), primers to amplify the upstream (7,8) and downstream (11,12) regions of the galK gene of 83972, with overlap to PA1_GFP, and primers to amplify GFP with the PA1 promoter (9,10).

*Bacterial cultures*
Single colonies were grown from frozen stocks on Luria broth agar plates at 37°C for 24h. Liquid cultures were prepared by inoculating 2 mL LB (Miller) broth (Carl Roth, Germany) using a single colony, and incubated overnight in a shaken incubator (37°C at 180 rpm). After

the overnight incubation, the cultures were diluted in 10 mL of fresh LB (Miller) and further incubated at 37°C for at least 6h. The cultures were then mixed in a desired proportion (approx. 1:1) according to their optical densities (OD600), with the exception of the experiment in Fig. 4, for which 10 ml overnight cultures were first centrifuged at 4000 rpm for 2 min, re-suspended in 1 mL to concentrate them 10x, and finally mixed in a 1:10 ratio (GFP:RIF$^S$ to mKate:RIF$^R$).

*Growth media and flow control*

We used LB Broth (Miller) sterilized by autoclaving at 121°C for 15 min, and supplemented with rifampicin (RIF) (Merck KGaA, Germany, concentrations as in the main text) for the experiments in Fig. 4. The medium was delivered by syringe pumps using plastic syringes (BD, USA). The syringes were connected to the microfluidic devices with PTFE tubing (Bola Bohlender, Germany, I.D. = 0.5 mm, O.D. = 1.0 mm), with identical lengths for both outlets to ensure equal hydraulic resistance. 0.5 mm OD needles were used to connect syringes to the tubing. Syringe pump PHD2000 (Harvard Apparatus, USA) was used for experiments in Figs. 1-2. For Fig. 4 we used a SyringeONE Programmable Syringe Pump (Darwin Microfluidics).

*Biofilm experiments*

For experiments in Figs. 1-2, the microfluidic device was flushed with 70% ethanol using a syringe mounted on the syringe pump, and left for 10 minutes. The syringe was then replaced with a syringe filled with LB medium and the microfluidic device was flushed with LB. Next, the syringe was replaced with a syringe filled with a bacterial suspension. The suspension was pumped through the device and left at room temperature for 30 minutes. Finally, the syringe was replaced with an LB medium syringe, and the device was placed on the XY microscope stage for imaging. The flow rate was set to 50 µl/h for the initial 16 h and changed to 200 µl/h afterwards.

For experiments in Fig. 4, the microfluidic device was prepared by first flushing it with a solution of 5% sodium hydroxide (Carl Roth, Germany) in 70% ethanol using a syringe mounted in a syringe pump, at a rate of 2 ml/h for 8 min. Afterwards, the device was flushed with 70% ethanol, followed by LB medium. A dense bacterial suspension was then introduced into the device using a syringe pump at a flow rate of 3 ml/h. When bacteria showed up in the main channel, the flow rate was reduced to 1 mL/h and periodically turned on and off for about 30 min, encouraging the bacteria to attach, while continuously imaging the wells until they contained hundreds of cells/well. The bacterial syringe was then replaced with an LB syringe. The flow rate was set to 50 µL/h for 500 µL (10 h), followed by an alternating fast/slow flow of 3 ml/h for 10 µL and 50 µl/h for 16 µL to reduce clogging of the main channel in the 180h-long experiment. Throughout the experiment, the medium was replaced with LB+RIF at 0.5 ug/ml, LB+RIF at 1 ug/ml, and pure LB as described in the main text.

We ran all experiments at room temperature (24-26°C); this helped to limit biofilm growth in the main channel.

*Microscopy*

Images were acquired using two fully automated Nikon Eclipse Ti2-E epi-fluorescent microscopes with automated XY stages and the Perfect Focus System, and controlled by MicroManager (7). One microscope used an ORCA-spark Digital CMOS camera C11440-36U (Hamamatsu, Japan), the other one an Andor Zyla 4.2 SCMOS camera (Oxford Instruments, UK). Depending on the experiment, we used two different objectives (20x and 40x). To acquire fluorescent images of mKate and GFP strains, we used filters with excitation/detection wavelengths of 532 – 554 nm/ 573 – 613 nm, and 457.5 – 487.5 nm/ 502.5 – 537.5 nm.

*Image and data analysis*
To analyze the data, we utilized custom Python and Mathematica® code to load and process the TIFF images generated by MicroManager. The code allowed us to perform the necessary data analysis and create plots for visualization. The code is available on GitHub as Jupyter and Mathematica notebooks (1).

*Figure 1.* To obtain the mean sector size, we calculated the autocorrelation function of pixel brightness along the cut through the middle of the well at ½ of the distance from the opening to the bottom surface. We then took the position of the first minimum of the autocorrelation function and used it to represent the mean sector size. This method offered several advantages compared to counting sectors in a thresholded image: (i) it was not sensitive to the absolute value of fluorescence signal, which could differ between wells and changed over time due to variations in RFP expression, (ii) it was not sensitive to minor variations in pixel intensity for neighboring pixels, thus avoiding sector over-counting caused by very narrow darker or lighter "streaks" within larger sectors. Moreover, the algorithm (iii) correctly predicted the average size of stripes for synthetic data with interleaved bright and dark stripes, (iv) it offered a more objective approach than manual sector counting.

*Figure 2.* We imaged the biofilm every second for about 3 minutes using the 40x objective. The biofilm grew only minimally during this time, and individual cells moved less than a pixel per time frame. Inspired by the methods shown before (8, 9), we determined the velocity field $\vec{v}(x, y)$ from subtle changes in pixel brightness caused by the biofilm's local motion. This method did not require tracer particles or feature detection within the biofilm. Briefly, we assumed that the flow in the biofilm caused the pixel intensity field $I(x, y, t)$ to evolve during a small time interval $dt$ as follows: $I(x, y, t + dt) = I(x, y, t) + dt\, \vec{v}(x, y) \cdot \vec{\nabla} I(x, y, t)$, where $\vec{\nabla}$ represents the (discrete) gradient operator. This linear set of equations (one for each pixel) could be solved numerically for $\vec{v}(x, y)$ using pixel intensities at two time points. Since the system of equations was underdetermined, we binned the image 32x in each direction and solved for the two components of the average velocity field within each 32x32 block of pixels using the least squares method. Additionally, we automatically selected the time separation $dt$ for each bin that yields the most accurate estimate for $v(x, y)$ at that location without violating the assumption of small changes ($v\, dt < 0.1$ bin size).

*Figure 4.* We first determined the intensity profiles in red (mKate) and green (GFP) channels along a horizontal line cutting through the well at ½ of its height, for all time points and wells. Pixel intensities in both channels were rescaled by the minimum and maximum values from the entire acquisition (all wells and time points), and then logarithmized as follows: $I_{\text{rescaled}} = \ln(I_{\text{raw}} - I_{\min} + 1) / \ln(I_{\max} - I_{\min})$. The 1d array obtained in this way was convolved with a top hat function of width 10 pixels to reduce noise. Next, we detected the boundaries between red and green sectors based on pixel intensity in each channel. A pixel was classified as belonging to a "red" sector if its intensity was higher in the red channel than in the green channel, and vice-versa for the "green" sector. Using sectors boundaries determined in this way, we calculated the number of sectors and their sizes for each well and time point. To find the fraction of sensitive (green) strain, we calculated the proportion of "green" pixels ($I_{\text{G,rescaled}} > I_{\text{R,rescaled}}$) along the line of pixels. This method worked reliably once green and red bacteria separated into sectors, which is the reason for the apparent decrease of the mean sensitive fraction in Fig. 4C during the first part of the experiment (no RIF), despite the red strain having a small fitness disadvantage. We also confirmed that this method gave qualitatively similar results to the manual counting of sectors (Fig. S3).

*Doubling time of bacteria in the biofilm from Fig. 4*
We tracked the movement of easy-to-distinguish features (brighter spots and swirls) in fluorescence images of biofilms growing in flat-bottomed wells during the first phase of the experiment presented in Fig. 4 (pure LB, no RIF). If $y_t, y_{t+\Delta t}$ denote the distance of a feature from the bottom at times $t$ and $t + \Delta t$, then the growth rate $\alpha$ can be calculated as

$$\alpha = \frac{\ln\left(\frac{y_{t+\Delta t}}{y_t}\right)}{\Delta t}.$$

We calculated $\alpha$ for all flat-bottomed wells, using 2-3 traceable features per well. We obtained the average value $\alpha = 0.22 +/-0.01$ h$^{-1}$, corresponding to the doubling time ≈3 h. We did not observe any significant correlation between $\alpha$ and $y_t$ of the feature (Pearson correlation coefficient $r = 0.10$, p-value $0.40$), which confirmed that growth did not depend on the depth (distance from the outlet) in the biofilm.
The growth rate was significantly lower in the biofilm from Fig. 4 than in the liquid culture (Fig. S5) at the same temperature. It was also lower than the growth rate from Fig. 2, in which the biofilm was less dense. Interestingly, the growth rate of single bacteria that occasionally attached to the surface of the main channel was very close to that of Fig. S5. Therefore, even if the growth medium flowing through the device was partially metabolised by bacteria living upstream, this was not significant enough to affect the growth rate.
In contrast to previous work (10), where mechanical confinement to a monolayer of cells has been suggested as a possible explanation of reduced growth rate, in our experiment the movement of cells is not constrained enough to cause axial compression, which could affect the growth rate (11). We hypothesize that cells in the biofilm switch to a slower-growing phenotype, perhaps as the effect of quorum-sensing (12).

*Relative fitness from sector expansion*
We used data from Fig. 4C to obtain the relative fitness $W_{\mathrm{G/R}}$ of the sensitive green strain compared to the resistant red strain. We first selected flat-bottomed wells in which the initial fraction of resistant (red) cells was between 0.1 and 0.4 at the beginning of the 45 h-long 1 μg/ml RIF phase from Fig. 4C. The [0.1, 0.4] range was chosen because the sector detection algorithm performed well in this range. Figure S4A shows plots of the resistant fraction versus time for the selected wells. We then ran the computer model in flat-bottomed wells for different resistant initial fractions and different relative fitness values $W_{\mathrm{G/R}}$. We assumed the doubling time of 3 h (see the preceding section) and the same duration of the simulation as the 1 μg/ml RIF phase. We selected runs for which the initial fraction was within $\pm 0.01$ of the experimentally determined values, thus the simulated curves (Fig. S4B) had the same initial distribution of resistant fractions as the experimental curves. Finally, we compared the theoretical and simulated average resistant fraction versus time curves obtained for different $W_{\mathrm{G/R}}$, and determined that the best fit was given by the model assuming $W_{\mathrm{G/R}} = 0.2$ and no adhesion.

*Relative fitness from liquid culture growth*

We used the method from previous work (13) to determine the growth rate of each strain at different concentrations of RIF. Briefly, we incubated bacteria in LB with different concentrations of RIF (0, 1, 1.5, 2, 2.5, and 3 μg/ml) in a 96 well micro-plate (200 μl/well) inside a plate reader, starting from two different initial cell densities, $N_0$ in rows A-D and $N_0/10$ in rows E-H. The $N_0$ dilution was an exponentially growing culture of OD600 ≈ $0.1$. We used one column of the 96 well plate for each RIF concentration. The plate reader (BMG LABTECH FLUOstar Optima) was set to incubate the plate at 25 °C in a cold room (4 °C) for enhanced temperature stability, with orbital shaking at 200 rpm for 10 s prior to OD measurement.

We measured the optical density (OD) of each culture every 2 min to obtain growth curves for approx. two days of incubation time. The exponential growth rate was then determined from the time shift between the growth curves for initial bacterial concentrations $N_0$ and $N_0/10$ .

Figure S5 shows that the two fluorescently-marked strains have almost identical growth rates in the absence of RIF. The sensitive green strain grows about 10% faster without RIF ($W_{\mathrm{G/R}} = 1.1$), but at 1 μg/ml used in experiments from Fig. 4 its relative fitness is $W_{\mathrm{G/R}} \approx 0.6$ compared to the red resistant strain. This differs from the estimate from the previous section based on fitting the computer model to the sector expansion experiment in the biofilm. We attribute this to a slower growth rate in the biofilm than in the liquid culture, which makes the $RIF^S$ strain more sensitive to RIF in the biofilm. Indeed, slower growth correlates with increased susceptibility: the minimum inhibitory concentration of RIF for the sensitive GFP strain is 16 μg/ml at 37 °C (fastest growth), 8 μg/ml at 30 °C and 2 μg/ml at room temperature (slowest growth).

*Relative fitness from a competition assay in bulk cultures*

The relative fitness of the *mKate*-labelled 83972 strain as compared to the wild type was determined by competing the strains against each other in LB broth, shaking at room temperature (25 °C). Overnight cultures of both strains were mixed in a 1:1 ratio, diluted 1000-fold into four replicate 10 mL cultures, and incubated until they reached the stationary phase. The ratio of the strains before and after competition was determined by plating $10^6$ dilutions of the initial mixture and of each of the end cultures on LB agar, and the number of red fluorescent and non-fluorescent CFUs was counted. The relative fitness (W) of the *mKate* strain was then calculated by the following formula (14), where $R_0$ is the frequency of *mKate* before the competition, $R_1$ is the frequency of *mKate* after the competition, and $F$ is the fold increase of bacteria during the competition as determined by the dilution:

$$W = \frac{\ln\left(\frac{R_1 \times F}{R_0}\right)}{\ln\left(\frac{(1-R_1) \times F}{1-R_0}\right)}.$$

We obtained $W = 0.97 \pm 0.01$, i.e., a very small fitness disadvantage of the mKate-labelled strain. For the purpose of our biofilm experiment, in which selection is very strongly suppressed as explained in the main text, we can consider both variants to have the same fitness.

### Supplementary Figures

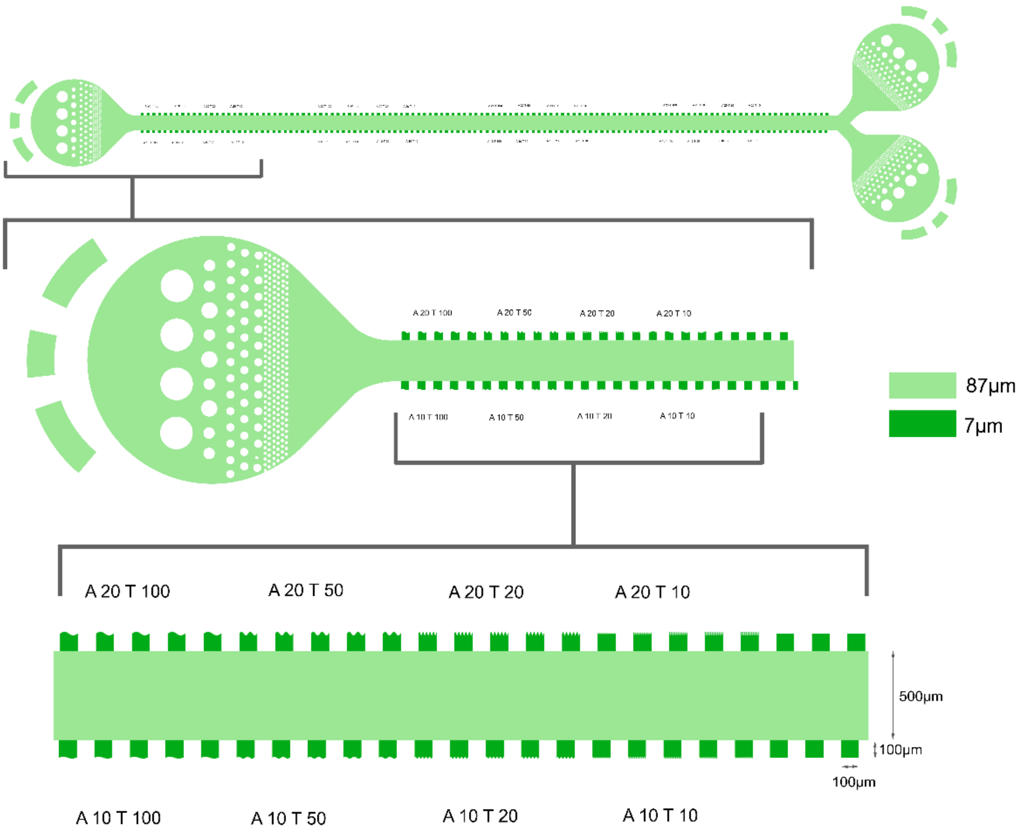


**Fig. S1. Design of the microfluidics device.** Top panel: the entire device. Middle and bottom panels: small sections of the main channel (width = 500 µm) with 100x100 µm wells of different types (corrugation period and amplitude) visible on both sides.

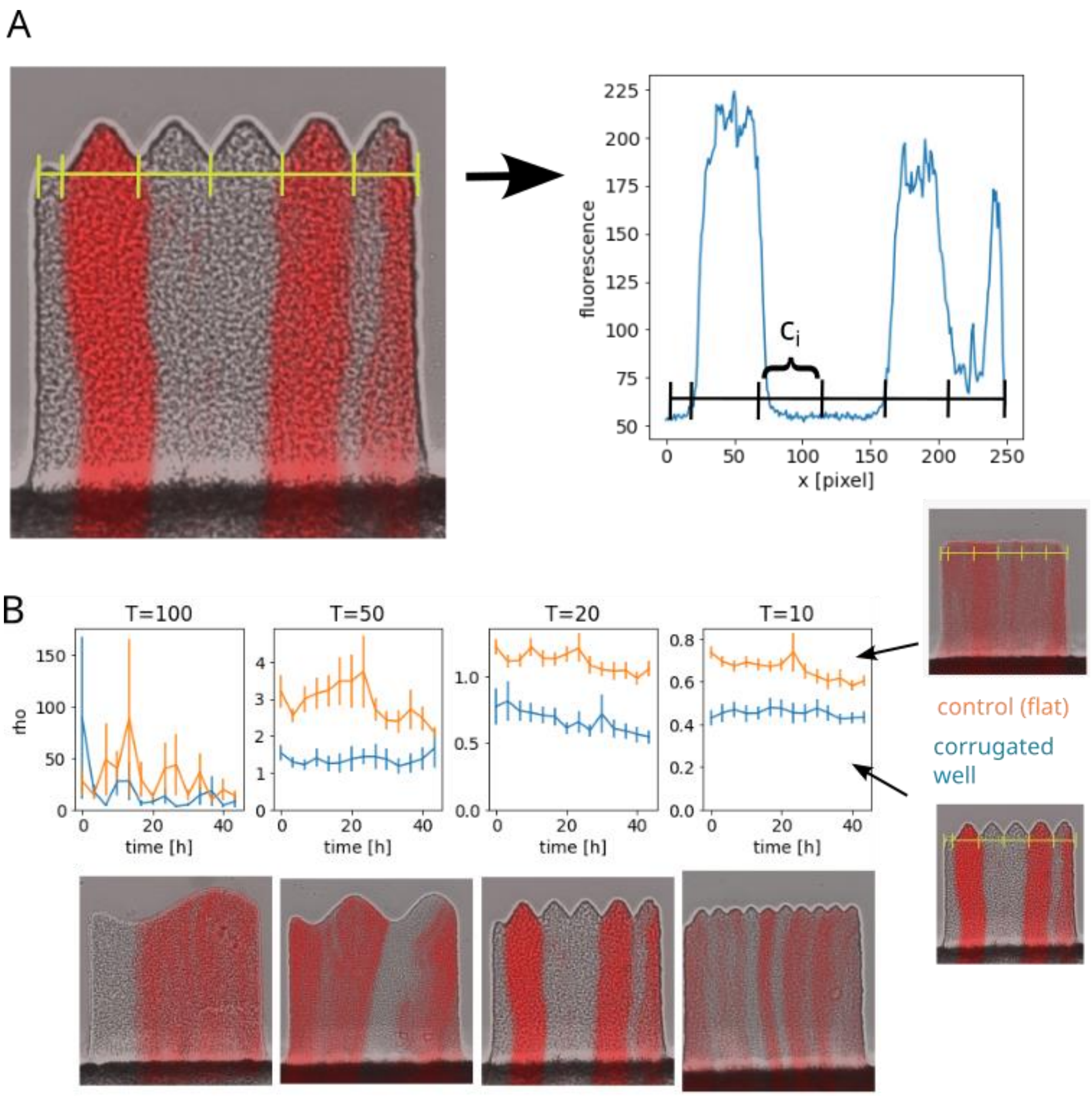


**Fig. S2. Intra- versus inter-well heterogeneity. (A)** Illustration of the method with which fluorescence heterogeneity is quantified. $\boldsymbol{c}_i$ is the vector of pixel intensities in the $i$th "pocket", calculated along the line (yellow) just above the undulations. The diversity ratio $\rho = \mathrm{E}(\mathrm{D}(c_1), \ldots, \mathrm{D}(c_N))/\mathrm{D}(\mathrm{E}(c_1), \ldots, \mathrm{E}(c_N))$ where $\mathrm{E}$ and $\mathrm{D}$ denote mean and standard deviation, respectively. **(B)** Diversity ratio $\rho$ versus time for wells with different undulation period $T$.

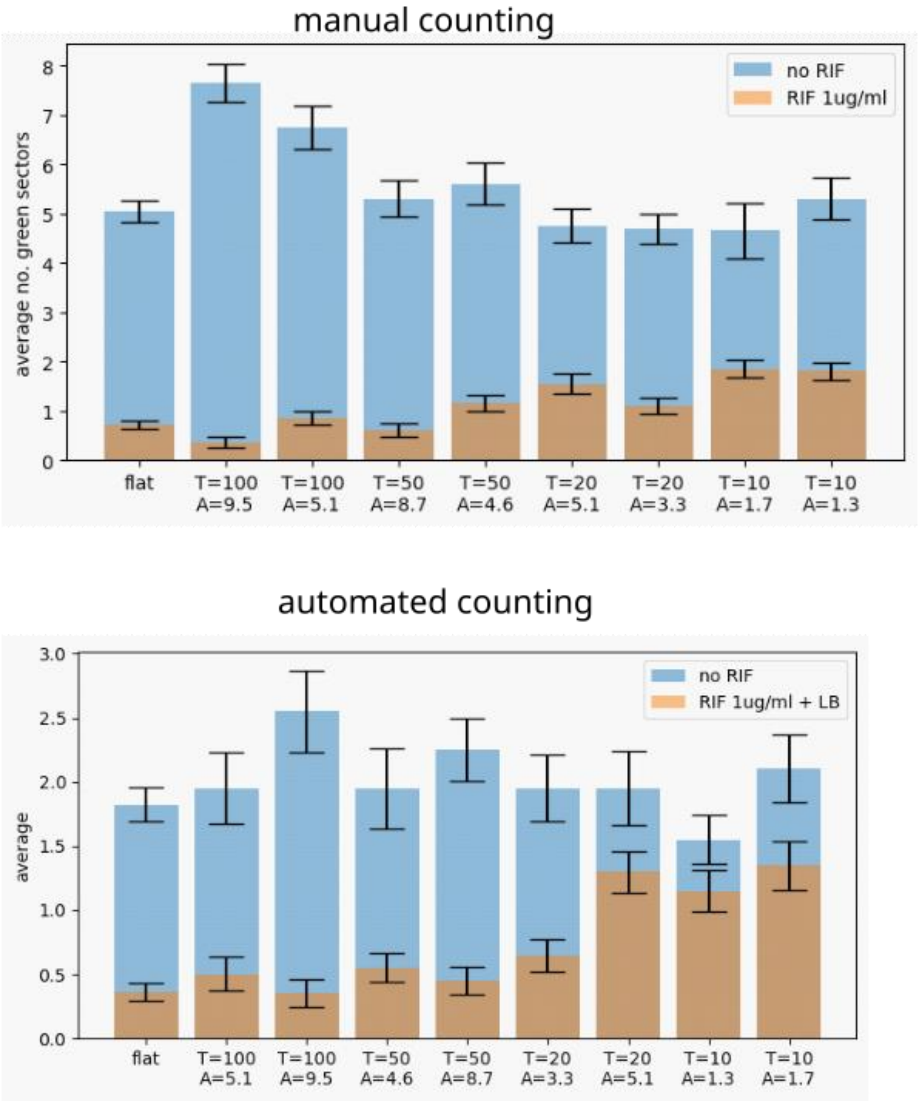


**Fig. S3. Average number of green sectors in the experiment from Fig. 4**. Upper panel: sectors counted manually, lower panel: sectors counted by a computer algorithm. The automated algorithm deliberately ignores very small sectors, thus the numbers are generally lower than with manual counting. However, both methods show the same trend: the ratio of the number of sectors after and before the RIF exposure increases with decreasing $T$.

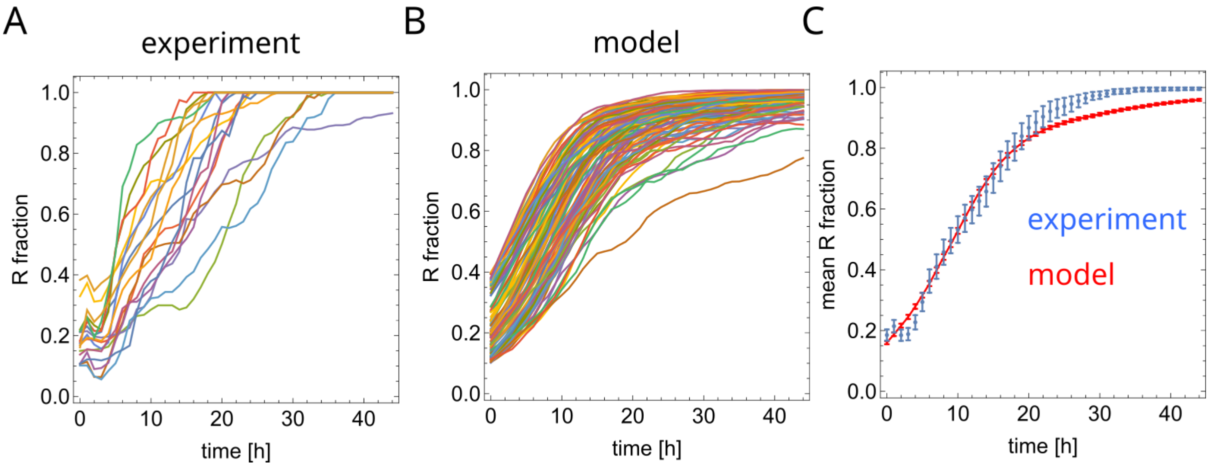


**Fig. S4. Relative fitness from sector expansion. (A)** The fraction of fitter (resistant) cells versus time in flat-bottomed wells (colors = different wells) during the 1 μg/ml RIF exposure experiment from Fig. 4C. Only wells in which the initial fraction at $t = 0$ h is between 0.1 and 0.4 have been selected. **(B)** The same fraction of fitter cells from the computer model, for the relative fitness of green (sensitive) to red (resistant) cells $W_{\mathrm{G/R}} = 0.2$, initial fraction of fitter cells having the same distribution as in panel (A) and the doubling time of 3 h. **(C)** Mean fraction of resistant cells for the experiment and the model, for $W_{\mathrm{G/R}} = 0.2$, which gives the best fit to the experimental data.

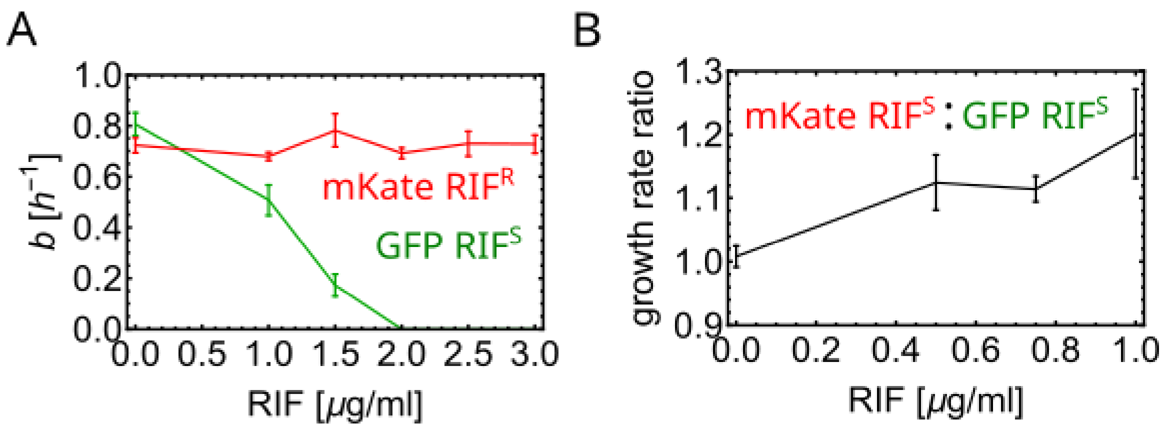


**Fig. S5**. **(A)** Growth rates of the RIF-sensitive GFP strain, and the RIF-resistant mKate strain. In the absence of RIF, the GFP strain grows ≈10% faster compared to the mKate strain. **(B)** The calculated ratio of the growth rates of mKate and GFP sensitive strains is consistent with no fitness difference in the absence of RIF, and a small growth advantage of mKate strain for non-zero RIF concentrations.

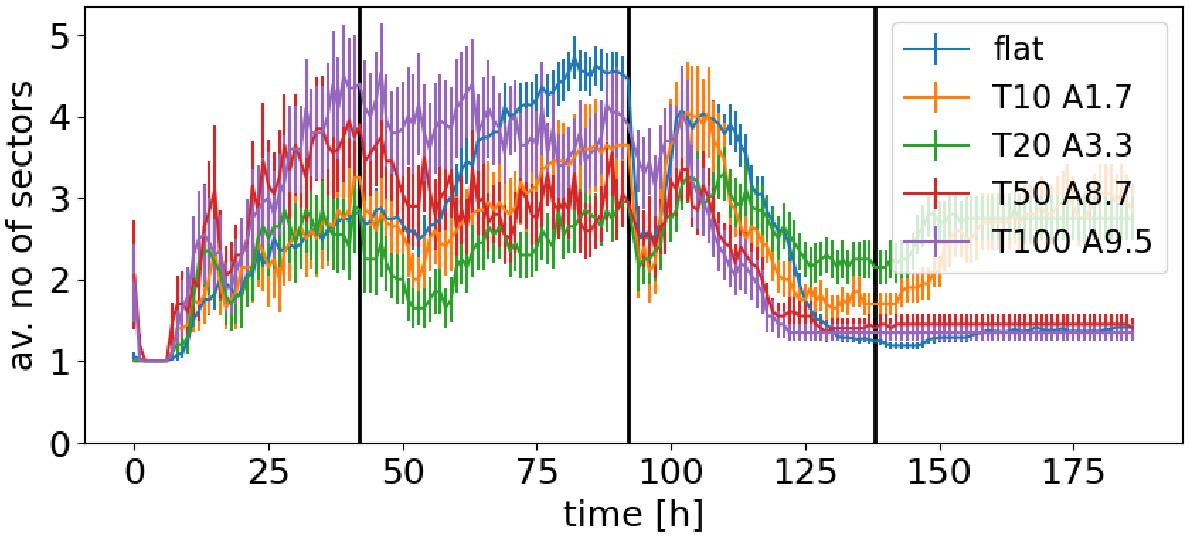


**Fig. S6. Average number of sectors (red + green) versus time, for the experiment from Fig. 4**. The sectors have been counted using the same automated algorithm as in Fig. 4.
